# Rigosertib Reverses Hypertrophic Cardiomyopathy in *RAF1*-Associated Noonan Syndrome

**DOI:** 10.1101/2024.11.25.623865

**Authors:** Levi Legler, Bing Xu, Tara Keshavarz Shirazi, Sereene Kurzum, Katya Marchetti, Chase Kessinger, Izabella Vredenburg, Yan Sun, Frank A. Dinenno, Damian Bohler, Angelika G. Aleman, Nelson A. Rodriguez, Simon Ng, Sophie Gao, Angela Wang, Mayte Suarez-Farinas, Hung-Mo Lin, Tirtha Das, Karen Ocorr, Ross L. Cagan, Bruce D. Gelb, Maria I. Kontaridis

## Abstract

**Background:** RASopathies constitute a group of rare genetic disorders caused by mutations in genes that reside along the canonical Ras/MAPK signaling pathway, affecting cell growth and differentiation. These syndromes, which include conditions like Noonan syndrome (NS), are characterized by developmental delays, distinctive facial dysmorphia, and a variety of cardiac defects, notably hypertrophic cardiomyopathy (HCM). Despite their prevalence and impact, therapeutic options for RASopathies remain limited. Rigosertib, a novel dual Ras/MAPK and PI3K/AKT pathway inhibitor, is currently in clinical trials for treatment of melanoma and recessive dystrophic epidermolysis bullosa. Here, we identify rigosertib as a candidate therapy for RAF1-associated HCM.

**Methods and Results:** Our Drosophila screen of clinically relevant drugs and compounds identified rigosertib as broadly effective across a panel of transgenic RASopathy fly transgenic models, indicating that rigosertib may be effective against multiple disease isoforms. Analysis of a Drosophila model targeting a *RAF1^L613V^* transgene to the heart found that rigosertib reduced aspects of cardiac hypertrophy. Rigosertib treatment prevented or regressed cellular hypertrophy in human induced pluripotent stem cell-(iPSC-) derived cardiomyocytes homozygous for the NS-associated *RAF1^S257L^* allele. We extended these findings to a mammalian model, using *Raf1^L613V/+^* KI mice to explore the therapeutic implications of rigosertib on RAF1-driven HCM. Longitudinal six-week treatment with rigosertib in these mice resulted in significant improvement in left ventricular chamber dimension and posterior wall thickness, total heart mass, size of individual cardiomyocytes (CMs), as well as reversal of cardiac hypertrophy. Rigosertib treatment also led to normalized fetal gene expression and inhibition of ERK and AKT pathway activities in primary CMs isolated from *Raf1^L613V/+^* mice. Cardiac function, as assessed by echocardiography, showed significant improvement in ejection fraction and fractional shortening, with molecular studies confirming downregulation of hypertrophic markers and signaling pathways. Together with the Drosophila data, these mammalian results support the potential and use for rigosertib to reverse pathological hypertrophy in NS through targeted pathway inhibition in patients. Moreover, in addition to its effects in the heart, rigosertib treatment in mice also significantly improved other NS-associated syndromic features, including increasing bone growth and correcting craniofacial abnormalities.

**Conclusions:** Taken together, our findings suggest rigosertib effectively normalizes and reverses RASopathy-associated HCM as well as other NS-associated syndromic features, supporting its potential for development as a promising treatment for RAF1-associated HCM and, potentially, other RASopathies-dependent pathologies. This study not only highlights the therapeutic potential of rigosertib but also demonstrates the utility of an integrated approach using Drosophila, iPSC and mammalian models to elucidate drug effects across complex biological systems.

## Introduction

RASopathies are a group of rare, nearly always autosomal dominant genetic disorders caused by pathogenic variants in members of the canonical RAS/mitogen-activated protein kinase (MAPK) signaling pathway [1]. RASopathies affect ∼1 in 1000-2000 live births, with the most common being Noonan syndrome (NS) [2]. The typical characteristics of NS include short stature, dysmorphic facial features, intellectual delays and disabilities, and cardiovascular issues; the latter include congenital heart disease (CHD)—most prevalently pulmonary valve stenosis—and hypertrophic cardiomyopathy (HCM) [2]. Indeed, HCM is the second most common cardiovascular manifestation in RASopathies, with a prevalence of 20-25%, exhibiting unique features that include early onset, rapid progression with congestive heart failure, and co-occurrence of CHD, leading to significant mortality [3].

NS is caused by gain-of-function (GOF) variants in multiple genes that affect RAS/MAPK signaling, including *PTPN11, SOS1, RAF1,* and *RIT1* genes, although pathogenic variants in other genes have also been described [4, 5]. There are several other genetic traits with overlapping clinical characteristics, such as Costello syndrome (CS), Noonan syndrome with multiple lentigines (NSML), and cardiofaciocutaneous syndrome (CFC). Like NS, these traits are caused by pathogenic alleles—typically GOF—in RAS/MAPK-related genes. NSML is the outlier, as it is most commonly associated with loss-of-function or dominant-negative mutations in *PTPN11* [6–9].

NS exhibits several well-established genotype-phenotype associations [4]. For example, *PTPN11*-associated NS, accounting for greater than 50% of all cases of NS [10], has an HCM prevalence of only 6%, significantly lower than the overall frequency of HCM in other NS-causing genes [3]. In this regard, the prevalences of HCM in *RAF1*-and *RIT1-* associated NS are 65% and 36%, respectively. While these two genetic forms of NS account for smaller percentages of NS pathogenesis (3-17% and 5-9%, respectively), they account for nearly 50% of HCM in NS and an even higher proportion of cases with early onset of congestive heart failure [3, 11]. These observations emphasize that different NS genes have distinct pathological effects and highlight the need for selective therapeutic approaches to treating NS-associated HCM.

Knock-in (KI) mice heterozygous for the NS-associated *Raf1 L613V* allele (*Raf1^L613V/+^*) develop HCM, with upregulated cardiac fetal gene expression and left-ventricular decompensation in response to pressure overload [12]. Agonist-evoked MAPK/extracellular signal-regulated kinase (ERK) activation is increased in *Raf1^L613V/+^*cardiomyocytes (CMs) and fibroblasts, suggesting it may play a critical role in growth factor-dependent modulation of ERK1/2-independent pathways [11–13]. RAS pathway inhibition, using the ATP-noncompetitive MAPK kinase (MEK) inhibitor mirdametinib, prevents HCM in *Raf1*-mutant mice [12], but only when treatment is initiated before the onset of hypertrophy. Based on this pre-clinical result, a limited number of infants and children with NS with severe HCM have been treated with the MEK inhibitor trametinib on a compassionate-care basis, with some evidence of regression of cardiac hypertrophy [14–16]. Clinical trial results are still needed to demonstrate formally the efficacy of this approach. Experience to date has shown some adverse effects of trametinib, particularly dermatologic, which can be limiting, and that not all affected patients respond to treatment [14, 15]. Thus, there is a need to identify other therapies that are efficacious and, ideally, less morbid, for NS-associated HCM.

To find additional small-molecule therapies with potential efficacy for the RASopathies, we developed a pre-clinical drug repurposing pipeline relying upon transgenic Drosophila models and identified that rigosertib could rescue pupal lethality in a *RAF1-*mutant RASopathy fly model [17]. Rigosertib (ON-01910 sodium salt) is a synthetic benzyl styryl sulfone with novel activities as a dual Ras/MAPK and phosphatidylinositol 3-kinase (PI3K)/AKT inhibitor [18, 19]. The primary mechanism of action for rigosertib remains unclear, with evidence supporting direct RAS pathway inhibition [20, 21] while other data indicate it acts as an inhibitor of microtubules [22–24]. Based on phase 1/2 clinical oncologic trials, rigosertib is well tolerated, with few side effects, the commonest being hemorrhagic cystitis, particularly in older men [25]. Of note, the range of side effects typically associated with anti-cancer agents that inhibit microtubules, such as peripheral neuropathies, has not been observed in trials of rigosertib. A Phase 3 clinical trial of rigosertib for myelodysplastic syndrome was unsuccessful [26]. Additional oncologic trials using rigosertib, alone or in combination with other agents, are ongoing or recently completed [27, 28]; it is also being trialed for recessive dystrophic epidermolysis bullosa-associated squamous cell carcinoma (Clinicaltrials.gov: NCT03786237 and 04177498).

In this study, we used a large panel of Drosophila RASopathy models to identify rigosertib as acting more broadly than other RAS pathway inhibitors, notably the MEK inhibitor trametinib. We demonstrated its efficacy in rescuing the cardiac hypertrophic phenotype in *RAF1* S257L fly hearts and *RAF1* S257L induced pluripotent stem cell (iPSC)-derived CMs. Finally, we undertook a trial of rigosertib in reversing the HCM phenotype in *Raf1^L613V/+^* mice, demonstrating that the drug both normalized cardiac status and restored linear growth and craniofacial anomalies. Our data support the view that rigosertib is a strong candidate as a highly effective, well-tolerated, and specific treatment for HCM in RASopathies, particularly for *RAF1*-associated HCM, as well as a potentially effective therapy for treating other NS-associated phenotypic effects.

## Materials and Methods

### Testing efficacy of rigosertib using transgenic Drosophila RASopathy models

#### Drosophila testing

We obtained rigosertib for use in these studies from Traws Pharma (formerly Onconova Therapeutics). We assessed rigosertib and trametinib for therapeutic efficacy using a quantitative Drosophila survival assay, as previously reported [29–31]. Briefly, rigosertib was dissolved in water (200 mM stock), and trametinib was dissolved in DMSO (200 mM stock). Drug was diluted in molten (∼50-60 °C) enriched fly media and then aliquoted into 5-ml vials to obtain final drug concentrations of 1, 10 or 50 µM. Based on previous analyses, after larvae consume drug-containing media, the circulating concentration of drug is ∼200-fold lower than the media concentration.

We used the driver *765-GAL4* line at 27 °C and crossed flies to *UAS-RASopathy* transgenic fly lines representing 14 different RASopathies variants observed in patients. Each transgenic line was fed rigosertib or trametinib, mixed into the media. Water only and DMSO only were used as untreated controls for rigosertib and trametinib, respectively. The ratio of treated:untreated surviving pupae/adults (expressed as “% rescue”) was used to assess drug efficacy. Biological replicates using 4-5 vials were performed per experimental condition. Each vial contained between 30-80 developing embryos, and absolute numbers of surviving pupae were counted.

#### Heart-specific testing in Drosophila

Previously published data indicate that expressing a RASopathy-associated transgene specifically in the Drosophila heart can recapitulate aspects of HCM [32, 33]. We focused on transgenic flies with cardiac-specific expression of *hRAF1^L613V^*because the allele is strongly associated with HCM in NS and pupal lethality could be rescued by rigosertib. We used the cardiac-specific driver line *tin-GAL4* and crossed them to our UAS-*hRaf1^L613V^* flies (*tin>hRaf1^L613V^*). Flies were raised on standard fly media or media containing 100 µM rigosertib or trametinib for one week starting at enclosure. Fly heart phenotypes were assessed with live imaging or with semi-intact hearts fixed in diastole and then stained with mouse anti-collagen IV (pericardin) 1:100, and/or phalloidin-cy3 1:1000 as previously described [34, 35].

#### Statistics in Drosophila studies

Effects of rigosertib and trametinib treatment were assessed across 14 Drosophila RASopathy models using linear models with treatment group as a factor. Differences across groups were assessed via F-test and, if p < 0.05, three hypotheses were tested: treatment effect for treatment 1 [*i.e*., rigosertib vs. control for rigosertib (water only)], treatment 2 [*i.e*., trametinib vs. control for trametinib (DMSO only)], and the differences in the treatment effects of rigosertib vs. trametinib. Such hypotheses were tested via contrast and adjusted for multiple hypothesis using the Bonferroni correction (*i.e*., nominal p value < 0.05 corresponding to rejection of the null hypothesis).

### Testing the efficacy of rigosertib using RAF1^S257L^ using human iPSC-derived CMs

#### Generation of the RAF1^S257L^ iPSCs

An induced pluripotent stem cell (iPSC) reporter line expressing the atrial-specific sarcolipin (*SLN*) gene [36] was used to generate *RAF1* S257L mutant lines, using the clustered regularly interspaced short palindromic repeats (CRISPR) dual nickase technology. Two guide RNAs (gRNA) targeting human *RAF1* with low off-target possibilities were designed using the MIT Optimized CRISPR design software. A 100-base-pair (bp) single-stranded oligonucleotide donor (ssODN) template was designed for introducing the target point mutation (Integrated DNA Technologies). gRNAs were inserted into the pCas9D10A_GFP vector (AddGene), and insertions were confirmed through Sanger sequencing (Genewiz).

Briefly, *SLN* iPSCs were plated on a 6-well matrigel-coated plate and confluent cells were harvested using Accutase (Innovative Cell Technologies, Inc.). One million *SLN* iPSC cells were nucleofected with Nucleofector Solution (Lonza), 6 µL of 100 µM ssODN template and 2.5 µg of the Cas9 plasmid containing the gRNAs for the *RAF1* S257L mutation and a GFP reporter. Cells were then plated across two wells in 6-well plates. Forty-eight hours post nucleofection, GFP+ cells were isolated via fluorescence-activated cell sorting (FACS) and plated on mouse embryonic fibroblast (MEF)-coated plates. Once single colonies were visible after a week, they were manually picked and expanded to 24-well plates and subsequently to 96-well plates. Restriction fragment length polymorphism (RFLP) analysis was performed to screen 198 individual colonies for the gene modification. A 500-bp sequence flanking the target sequence was PCR amplified from DNA extracted from the iPSC colonies, and samples were digested with restriction enzyme *Sal*I overnight. Digested samples were resolved on 1% agarose electrophoresis gel, and 52 positive *RAF1^S257L^* clones were sequenced via Sanger sequencing to confirm the correct gene modification. One homozygous *RAF1^S257L^* modified clone was identified; this clone screened negative for off-target modification sites provided by the MIT design software. It was then purified through dilutional cloning, expanded and stored in -80 °C.

#### hiPSC differentiation

Embryoid bodies (EBs) were generated as described previously [37], with the following modifications: hiPSCs were cultured on Matrigel-coated plate for three days until they were 80-90% confluent. Cells were then dissociated using 0.6 mg/mL collagenase B (Roche) for 1 hour in a 37 °C incubator. Once the cells lifted off the plate, they were collected and washed with DMEM (Thermo Fisher Scientific), DNase (Millipore), thiazovivin (Millipore), 0.05% bovine serum albumin (Thermo Fisher Scientific) and centrifuged for 3 minutes at 1200 rpm. Cell pellets were resuspended in differentiation media containing RPMI 1640 medium (Thermo Fisher Scientific), B27 supplement plus insulin (Thermo Fisher Scientific), 1% PenStrep (Thermo Fisher Scientific), 2 mM GlutaMAX (Thermo Fisher Scientific), 50 ug/mL ascorbic acid (Sigma), 150 ug/mL transferrin (Sigma), and 50 ug/mL monothioglycerol (MTG) (Sigma). EBs were distributed in 6-well plates. In order to induce CM differentiation, the media contained 2 ng/mL BMP4 (R&D Systems) on Day 0, 20 ng/mL BMP4, 20 ng/mL activin A (R&D Systems) and 5 ng/mL bFGF (R&D Systems) on Day 1, and 10 µM IWP-2 (Tocris) and 5 ng/mL VEGF-A (R&D Systems) on Day 3 and stored at 37 °C in 5% CO_2_, 5% O_2_ and 90% N_2._ No changes were made between Days 3 to 5. After Day 5, the culture medium was changed every two days.

#### Flow cytometry

EBs were dissociated with 1 mg/mL collagenase II (Worthington Biochemical) in a 37 °C incubator for 1 hour, washed and centrifuged for 3 minutes at 1200 rpm. To further dissociate the EBs, the pellet was resuspended in 2 mL TrypLE (Thermo Fisher Scientific), kept in 37 °C water-bath for 5 minutes, washed, centrifuged, and resuspended in 10% FBS staining buffer. The sample was incubated with anti-human SIRP⍺-PE/CY7 (1:500) (Biolegend) and CD90-FITC (1:250) (Biolegend), incubated on ice for 1 hour, washed and resuspended in FACS buffer containing 2% fetal bovine serum in PBS, DNase (1:100), and thiazovivin (1:5000) dilution and DAPI. FACS-isolation was then performed on an Aria cell sorter (BD Biosciences).

#### Immunofluorescence

Single cells from FACS-isolation or whole EB dissociation were plated on Matrigel-coated plates. Once the cells were fully attached to the plate, they were fixed with 4% paraformaldehyde at room temperature for 15 minutes, washed with PBS, blocked with 0.5% saponin for 1 h and stained with primary antibody anti-human cardiac troponin T (cTNT) overnight. Cells were washed the next morning and stained with secondary antibody (Alexa 448) for 1 hour, washed again and stained with DAPI for 20 minutes prior to imaging. Images were taken on EVOS fluorescence microscope and analyzed via the ImageJ software.

#### Assaying iPSC-derived CM size

To assess CM size, EBs were dissociated, isolated through FACS, and FACS-isolated iPSC-derived CM and fibroblast-like cells (FLCs) were plated on matrigel coated plates. Control iPSC-derived CMs were either plated alone or with control FLCs, and *RAF1^S257L^* iPSC-derived CMs were either plated alone or with *RAF1^S257L^* FLCs, all at a density of 25,000 cells per well and a 60:40 CM to FLC ratio. Cells were fixed after 4 days, as described above, and the cellular area of cTNT^+^ cells was measured. For the time course experiment to determine the onset of cellular hypertrophy, CMs were fixed on Days 10, 15, 20, 23, and 25.

#### Rigosertib treatment

To determine whether the increased cell size observed in *RAF1^S257L^*iPSC-derived CMs could be prevented or reversed, we designed a serial dilution drug screen experiment with rigosertib. Rigosertib concentrations used were 10, 100, and 1000 nM. Rigosertib was dissolved in dimethyl sulfoxide (DMSO) for a final concentration of 1 uL/mL. DMSO (1 µL/mL) was used as a negative control. Whole EBs were dissociated with collagenase II and TrypLE as described above on Day 14. Prior to EB dissociation, 24-well plates were precoated with 60 µL of matrigel at the center of each well. Single cells were resuspended in 500 µL of RPMI medium and thiazovivin (1:5000), and 1 µL of the relevant rigosertib stock per well. Each drug concentration was tested in five wells of 24-well plates. Rigosertib treatment was from Day 15 to 19 for the prevention experiment and from Day 20 to 24 for the reversion experiment, and the media was changed every day. The cells were then fixed on Day 20 and 25 for the prevention and the reversion experiments, respectively, as described above, and cellular areas were measured. The person performing the imaging and measuring cell areas was blinded to the treatment condition.

#### Statistical analysis of iPSC-derived CMs

The cell areas of the rigosertib-treated *RAF1^S257L^* iPSC-derived CMs were compared to DMSO-treated *RAF1^S257L^* iPSC-derived CMs. The cell size ratio was calculated by dividing individual *RAF1^S257L^* iPSC-derived CM area by the average cellular area of the DMSO-treated mutant cells. Experiment analysis was performed using mixed-effects model for repeated measures. The least square means of the cell size ratio was estimated between different concentrations of rigosertib and DMSO control. Rigosertib and its three concentrations were set as fixed effects, and the wells of the cell-culture plates from which the cellular areas were measured were set as random effects. The significance levels were set as *P <* 0.05. The restricted maximum likelihood method with the Kenward-Roger correction of degrees of freedom was used to estimate the effects of rigosertib and its three concentrations on the cells. The null hypothesis was that the treatment, relative to DMSO control, did not reduce cell size (cell size ratio is 1) and the alternative hypothesis was that the cell size ratio was not 1. Statistical analyses were performed with SAS.

### Testing the efficacy of rigosertib in a KI Raf1^L613V/+^ mouse model

#### Mice

*Raf1^L613V/+^* (B6N;129S6-Raf1tm1.1Bgn/Mmucd, 036521-UCD) were obtained from the Mutant Mouse Resource & Research Centers (MMRRC) at The Jackson Laboratory, an NIH-funded strain repository, and were donated to the MMRRC by Benjamin G. Neel, M.D., Ph.D., New York University. Clippings were taken from the tip of the tails (1 mm) of 18–21-day old mice, DNA was extracted and subjected to PCR to assay for the *Raf1^L613V^* allele using the primers: sense 5’-ATC CCC TGA TCT CAG CAG GCT CTAC-3’; antisense 5’-AGT AGT CTA GGT CCT TAG CAG CAGC-3’, with subsequent restriction enzyme digestion using *Dra*III (New England Biolabs) to determine genotype. Animals were kept on a 12-hour light-dark cycle, at 22 °C and food was distributed *ad libitum*. Animals were maintained and utilized for the experiments herein by crossing *Raf1^L613V/+^* mice with mixed C57BL/6 x 129 wildtype (WT) mice (Jackson Labs). All animal procedures were approved by the Masonic Medical Research Institute Animal Care and Use Committee. MMRI’s PHS assurance number is D16-00144(A3228-01), and it is an AAALAC-accredited institution (#001865).

#### Treatment of mice with rigosertib

Rigosertib was provided by Onconova Therapeutics (now Traws Pharma). Rigosertib was dissolved in sterile filtered water at 0.01 mg/uL and injected intraperitoneally (IP) at 100 mg/kg body weight, twice daily, for either three or six weeks, as indicated. Prior to sacrifice, mice were fasted for 4 hours, following the final round of injections, to normalize signaling effects for subsequent molecular assessments.

#### Echocardiography

Transthoracic echocardiography and left ventricular functional assessments were conducted on non-anesthetized animals as described previously [8, 9], using a Visual Sonics Vevo 3100^®^ high-frequency ultrasound rodent imaging system. Briefly, mice were trained for three consecutive days prior to baseline recordings to acclimate to the echocardiography procedure. Baseline measurements were made at 8 weeks of age and were repeated at 11 weeks and 14 weeks of age, to assess effects of three and six weeks of rigosertib treatment, respectively. Hearts were imaged in the two-dimensional parasternal short-axis view, and an M-mode echocardiogram of the mid left ventricle was recorded at the level of papillary muscles. Heart rate, interventricular septum (IVS), left ventricular posterior wall thickness (LVPW), and end-diastolic and end-systolic internal diameters of the left ventricle (LVIDd and LVIDs, respectively) were measured from M-mode images.

#### Anatomical measurements

Body weights for each mouse were recorded weekly to determine dosing. At the time of sacrifice, mice were weighed and then measured from nose tip to tail base to determine body length. At harvest, the heart and lungs were removed, washed with PBS, blot dried and weighed. The right tibia of each mouse was also removed, and length measurements were taken.

#### Craniofacial morphometry

We utilized μCT scans using a Locus Ultra MicroCT Scanner (GE Healthcare) to measure changes in craniofacial morphometry. Three-dimensional images of the craniofacial skeleton were generated and analyzed with GEHC MicroView Software (GE Healthcare). Skull measurements were made according to the Standard Protocol and Procedures from The Jackson Laboratory (http://craniofacial.jax.org/standard_protocols.html).

#### Microscopy and histology

Hearts for morphometry and histochemistry were flushed with PBS and then fixed in 4% paraformaldehyde for 24 hours before paraffin embedding. Serial sections (5-μm thick) of heart tissue were taken in the transverse plane from each mouse at the level of the papillary muscle, and then stained with either reticulin or hematoxylin/eosin (H&E). Images of these sections were taken using the Keyence BZ-X810 fluorescence microscope at both 40x and 400x magnification, with an exposure of 1/60 s at 50% transmittance, and then labeled with a 50-mm measure bar.

Tissue sections of hearts were used to assess the cross-sectional length, width, and area of C, where centrally located nuclei (to ensure the same plane of sectioning) were measured using ImageJ 1.41 software (developed by Wayne Rasband; http://rsbweb.nih.gov/ij/). Three individual samples were analyzed for each genotype, with five different fields from four mice per group and three sections per heart. The total number of myocytes counted was 1-2 x 10^3^ cells per mouse.

#### Adult primary CM isolation

Mice were injected with intraperitoneal heparin (40 units/mouse), and their hearts were isolated and perfused via the aorta. The perfusion buffer consisted of KCl 14.7 mM, NaCl 120.4 mM, KH_2_PO_4_ 0.6 mM, Na_2_HPO_4_ 0.6 mM, MgSO_4_7H_2_O 1.2 mM, 2,3 butanedione monoxime 10 mM, taurine 30 mM, HEPES 10 mM, and glucose 5.5 mM. Hearts were digested with collagenase II digestion buffer (2 mg/ml) for approximately 8-10 minutes. Each heart was then cut from the cannula and placed in the dish with digestion buffer and stopping buffer [12.5 µM CaCl_2_ and 10% exosome-depleted FBS in perfusion buffer]. Isolated CMs were then cultured in 6-well plates (treated with laminin), with myocyte culture medium [ScienCell #6201], 5% exosome-depleted FBS, and 1% penicillin/streptomycin. For signaling experiments, isolated CMs were starved and either treated with vehicle (water) or rigosertib (10 nM) for 12 hours and then stimulated with either vehicle (DMSO), angiotensin II (20 ng/ml) or insulin-like growth factor (IGF) (20 ng/ml), as indicated, for 5 minutes.

### Biochemical studies associated with Raf1^L613V/+^ KI mice

#### Immunoblotting

Tissue or cell lysates were prepared by homogenizing the tissue or lysing the cells, respectively, in radioimmunoprecipitation (RIPA) buffer (25 mmol/l Tris-HCl [pH 7.4], 150 mmol/l NaCl, 0.1% SDS, 1% NP-40, 0.5% sodium deoxycholate, 5 mmol/l EDTA), 1 mmol/l sodium fluoride, 1 mmol/l sodium orthovanadate, and a protease cocktail at 4 °C, followed by sonication. The homogenate tissue was centrifuged for 10 minutes at 15000 x g at 4 °C, after which the supernatant was collected. Protein concentration was estimated using a “Pierce Rapid Gold BCA Protein Assay Kit” as directed by the manufacturer (Thermofisher).

For immunoblots, 30 μg of total protein lysates was resolved through a 10% polyacrylamide gel by sodium dodecyl-sulfate polyacrylamide gel electrophoresis (SDS-PAGE) and then transferred to a 0.2-µm nitrocellulose membrane. The resulting membranes were immunoblotted using antibodies against anti-AKT (#9272), anti-phospho-AKT (Ser473) (#9271), anti-ERK1/2 (#4695), and anti-phospho-ERK 1/2 (Thr202/Tyr204) (#9101) (Cell Signaling Technology). Visualization of the primary antibody binding was performed using the Li-Cor anti-rabbit 680 IRDye system in conjunction with the Li-Cor Odyssey FC. Quantification of secondary antibody fluorescence was performed using ImageStudio Lite Ver 5.2.

#### RT-qPCR

The apex of the left ventricle was utilized for extraction of total RNA using the RNeasy fibrous tissue kit (Qiagen). Reverse transcription of 1 µg of RNA was conducted using the iScript cDNA synthesis kit (BioRad). RT-qPCR samples were tested in triplicate. SYBR green (Applied biosystems) was used per manufacturer’s instructions. Primers targeting expression of fetal genes *Anp, Bnp, Myh6,* and *Myh7* were generated, and all levels of resulting expression were normalized to the average of two housekeeping genes *β-actin* and *18s ribosomal RNA (18S).* Data were quantified using the comparative C_T_ method (ΔΔCT). For primer sequences and PCR conditions, please see the supplemental information (**Table S1**).

### Statistical analyses

All values in graphs are expressed as the mean ± SEM. Normality was tested with the D’Agostino-Pearson normality test. If the data showed a normal distribution, pairwise testing was performed with the student’s t test or multiple group comparisons were performed by 1-or 2-way ANOVA, followed by post-hoc Bonferroni correction (GraphPad Prism 9). Significance level was set to 0.05.

## Results

### Rigosertib treatment showed efficacy against most RASopathy Drosophila models

Our goal was to identify a lead therapeutic compound effective against multiple RASopathy isoforms. Given the challenges of recruiting a sufficient number of patients with the individual RASopathy genes, we tested candidate drugs/compounds using 14 previously generated Drosophila models, each of which expressed a different human or Drosophila RASopathy gene variant under the control of an inducible UAS promoter [17]. In our screen of RAS relevant compound and drugs ([17]), rigosertib proved unique in its broad efficacy. When treated with rigosertib at three different doses (1, 10, and 50 µM) in the fly media, we observed significant pupal rescue for 8/14 (57%) of the RASopathy fly models (**Figure 1A**). By comparison, feeding flies the potent MEK inhibitor trametinib led to rescue of pupal lethality in 3/14 (21%) models; all three were also rescued by rigosertib (**Figure 1B**). For two RASopathy fly models—*KRAS^G12D^* and *PTPN11*^Y279C^—the treatment effect size rescue was statistically greater for rigosertib than trametinib; no fly model showed a greater effect size rescue by trametinib than rigosertib (**Figure 1B**). These data together suggest that rigosertib is especially effective in rescuing pupal lethality in Drosophila RASopathy models, outperforming the RASopathy candidate therapeutic trametinib.

**Figure 1.**
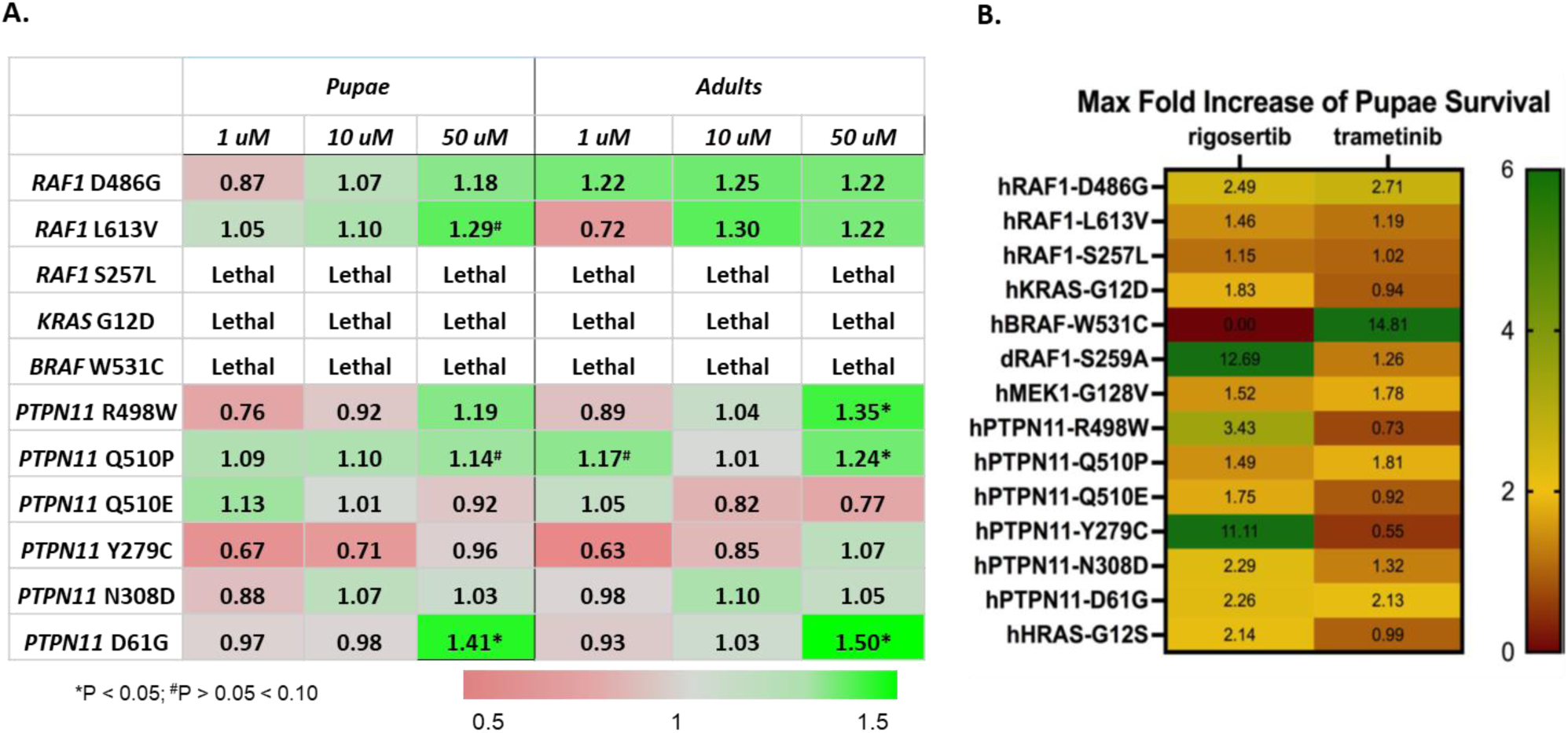
Rigosertib treatment shows efficacy against some, but not all, RASopathy drosophila models. **A.** The fold rescue of 1, 10, or 50 μM rigosertib treatment in RASopathies Drosophila models, as compared to untreated flies of the same genotype, for both pupae (left) and adults (right). Fold changes are indicated in the color code bar at the bottom of the table. Significance or trend to significance are indicated with * and ^#^, respectively. **B.** Maximal fold increase in pupal survival following treatment with 100 µM of either rigosertib or trametinib for one week starting at enclosure, as compared to untreated RASopathy Drosophila models. For pupal and adult viability analysis, mean and standard error of the mean (SEM) were calculated from 4-5 vials (biological replicates) per experimental condition. Each vial contained between 30-80 developing embryos, and absolute numbers of surviving pupae/adults were compared to diluent-only treatment to obtain % rescue.

### Rigosertib rescues the RAF1^L613V^ Drosophila HCM phenotype

*RAF1*-associated RASopathies are strongly associated with HCM, which is a significant source of morbidity and mortality in these patients. To assess the effects of rigosertib on cardiac pathology in Drosophila, we used the heart-specific *tin-Gal4* promoter to drive the UAS-human *RAF1^L613V^* transgene (*tin>hRAF1^L613V^*), resulting in expression of a cardiac-specific *RAF1^L613V^* NS allele, a variant strongly associated with HCM. We found that adult Drosophila hearts expressing *tin>hRAF1^L613V^* showed multiple deficits, including significant increases in heart systolic and diastolic diameters, and significant increased total heart tube are (Fig. A-C). The fibrous extracellular collagen IV (pericardin) network associated with the heart tube was also significantly increased (**Figure 2E-F**). In addition, optical sections through the conical chamber (the largest and most anterior portion of the heart tube) stained for F-actin showed significant hypertrophy of the myocardium in *tin>hAF1^L613V^*hearts (**Figure 2G-I**). These data suggest that the *tin>hRAF1^L613V^* fly model recapitulates key aspects of HCM observed in NS patients with inherited *RAF1* pathogenic alleles. Importantly, feeding adult flies with rigosertib for one-week post-eclosion significantly improved the heart systolic and diastolic diameters (**Fig. 2A-C**), fibrosis (**Fig. 2E, J-L**), and mean thickness (**Fig. F, 2J-L**), suggesting the potential for rigosertib to serve as an effective therapeutic agent for treating RAF1-associated HCM.

**Figure 2.**
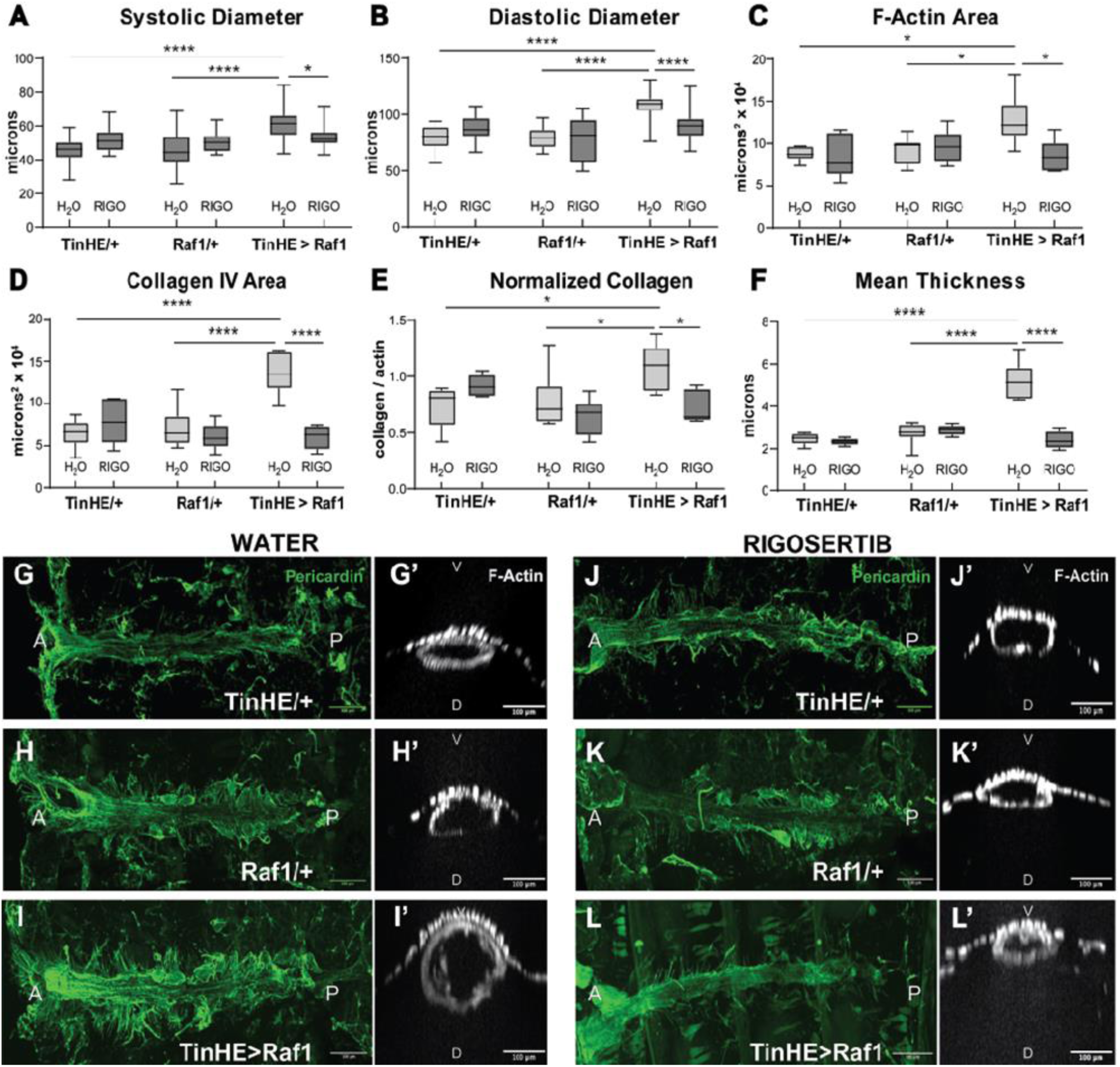
Rigosertib rescues the *RAF1^L613V^* Drosophila HCM phenotype. Using targeted heart-specific *tin-Gal4* promoter to drive the UAS-human *RAF1^L613V^*transgene (*tin>hRaf1^L613V^*), we observed significant changes in systolic diameter **(A)**, diastolic diameter **(B)**, F-actin area **(C)**, collagen IV area **(D**), normalized collagen levels **(E),** and mean thickness **(F),** indicative of pathological hypertrophy; these changes were reversed in the presence of rigosertib (Rigo) at the concentration of 100 µM in the fly media. Optical cross sections from Z stacks of pericardin-stained Drosophila hearts (**G-L**) and their F-actin-stained myocardial layer (ML) (**G’-L’**), which is significantly increased in the *hRAF1^L613V^*-expressing hearts. Hearts from flies fed 100 µM rigosertib for 1 week starting at eclosion showed significantly reduced thickness of the myocardial layer. In addition to thickening, hRAF1^L613V^ hearts displayed poorly aligned actin and collagen bundles. N= 3-4 experimental replicates. Data represent mean ± SEM; *p<0.01, ****p<0.0005, and all p values were derived from ANOVA with Bonferroni post-test when ANOVA was significant.

### Human iPSC derived RAF1^S257L^ CMs respond to rigosertib treatment

#### Generation and characterization of a RAF1^S257L^ iPSC line

We utilized an iPSC reporter line expressing the atrial-specific sarcolipin (SLN) gene [36] to generate a *RAF1* S257L mutant line, using the clustered regularly interspaced short palindromic repeats (CRISPR) dual nickase technology. The resultant *RAF1^S257L^* iPSC line was confirmed as homozygous by Sanger sequencing (**Figure 3A**). To assess hypertrophy in the context of *RAF1^S257L^*, iPSCs were differentiated to EBs, dissociated on Day 30, FACS-sorted and plated on 24-well plates. We used a time-course experiment to establish the timing of hypertrophy in these mutant cells. EBs were dissociated, and single cells were cultured on 24-well Matrigel-coated tissue culture plates. Cells were fixed and cellular area was measured at the following differentiation time points: Days 10, 15, 20, 23, and 25. Our results indicate that the *RAF^S257L^* iPSC-derived CMs display hypertrophy between Days 15 and 20 and that the cellular area of these mutant iPSC-derived CMs is significantly larger on Day 20 than in the control iPSC-derived CMs (p < 0.01) (**Figure 3B**).

**Figure 3.**
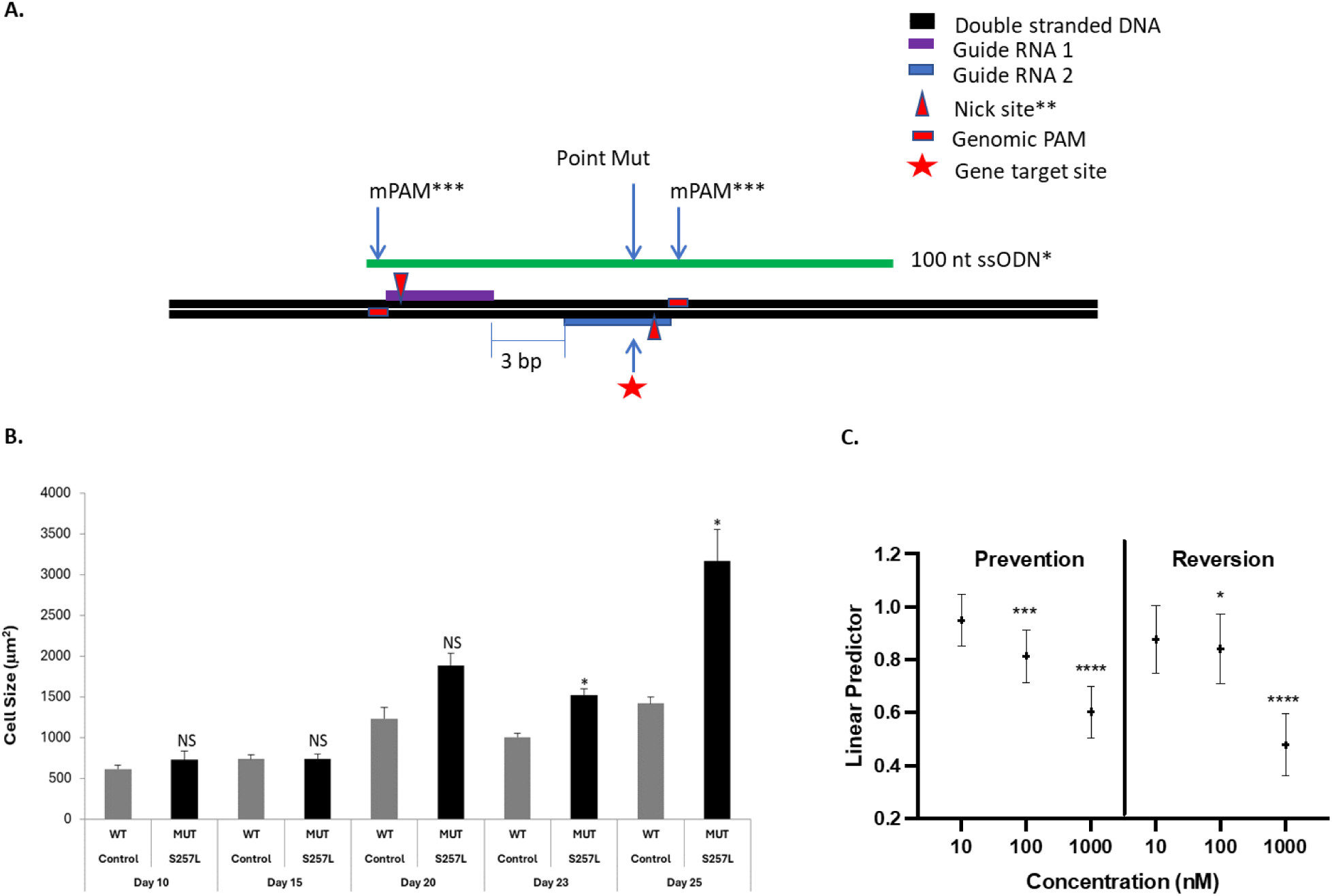
Human iPSC derived *RAF1^S257L^* CMs respond to rigosertib treatment. **A.** We utilized an iPSC reporter line expressing the atrial-specific sarcolipin (SLN) gene to generate a homozygous *RAF1* S257L mutant line using the clustered regularly interspaced short palindromic repeats (CRISPR) double nickase technology. *100 nt ssODN is sense to the coding strand with ∼50bp arms flanking the mutation site; **nicks introduced using Cas9n (D10A); ***silent mutations for mPAM sites. **B.** Control or *RAF^S257L^* iPSC-derived CMs were fixed, and cellular area was measured at the following differentiation time points: Days 10, 15, 20, 23, and 25. NS: Not Significant; * p<0.01. **C.** Homozygous *RAF^S257L^* iPSC-derived CMs were treated with rigosertib (10, 100, or 1000 µM) for five days to prevent cellular hypertrophy (treatment started on Day 20, left) or reverse existing cellular hypertrophy (treatment started on Day 25, right). Shown are the 95% confidence intervals for estimates of the least square means for cell areas for each treatment condition and statistical significance levels determined using mixed-effects modeling for repeated measures.

#### Rigosertib treatment

*RAF1^S257L^* iPSC-derived CMs were treated with rigosertib, either (i) prior to the onset of cellular hypertrophy (prevention) or (ii) after hypertrophy was already established (reversion). For the prevention study, treatment with rigosertib, initiated at Day 15 for five days, significantly reduced *RAF1^S257L^* iPSC-derived CM areas at 100 and 1000 µM (**Figure 3C**); treatment with 10 µM had no significant effect. For the reversion study, treatment with rigosertib was initiated at Day 20 and continued for five days; again, *RAF1^S257L^* iPSC-derived CMs showed a statistically significant reduction in cell area for dosing at 100 and 1000 µM (**Figure 3C**); no effect was observed at 10 µM. Of note, treating with 1000 µM rigosertib, for either prevention or reversion, resulted in cell areas similar to control iPSC-derived CMs. Rigosertib-treated iPSC-derived CMs remained healthy in culture and continued to show spontaneous beating patterns that were indistinguishable from untreated control and *RAF1^S257L^* iPSC-derived CMs.

### Rigosertib treatment of KI Raf1^L613V/+^ mice reversed hypertrophic cardiomyopathy

To validate the Drosophila and iPSC-derived CM findings and to better investigate the impact of rigosertib on mammalian HCM, we next examined the effects of rigosertib on *Raf1^L613V/+^* KI mice. As previously described, both male and female *Raf1^L613V/+^* mice developed cardiac hypertrophy by 8 weeks of age [12]. To assess the effects of rigosertib in these mice, we injected male or female 8-week-old mice intraperitoneally (IP) with either vehicle (water) or rigosertib at a dose of 100 mg/kg body weight, twice daily; we then assessed their cardiac physiological and functional parameters after six weeks of treatment. Our experimental dose was selected based on maximum tolerable dose (MTD) studies previously conducted at Traws Pharma (formerly Onconova Therapeutics), where 100 mg/kg/day was found to be both sub-MTD and efficacious in a long-term study [20]. No mortality occurred in our cohort as a consequence of treatment with rigosertib at this dose for this time period. We found that rigosertib reverted and completely normalized the HCM cardiac phenotype in both male and female *Raf1^L613V/+^* mice (**Figure 4A-B**, **Figure S1A-B**).

**Figure 4.**
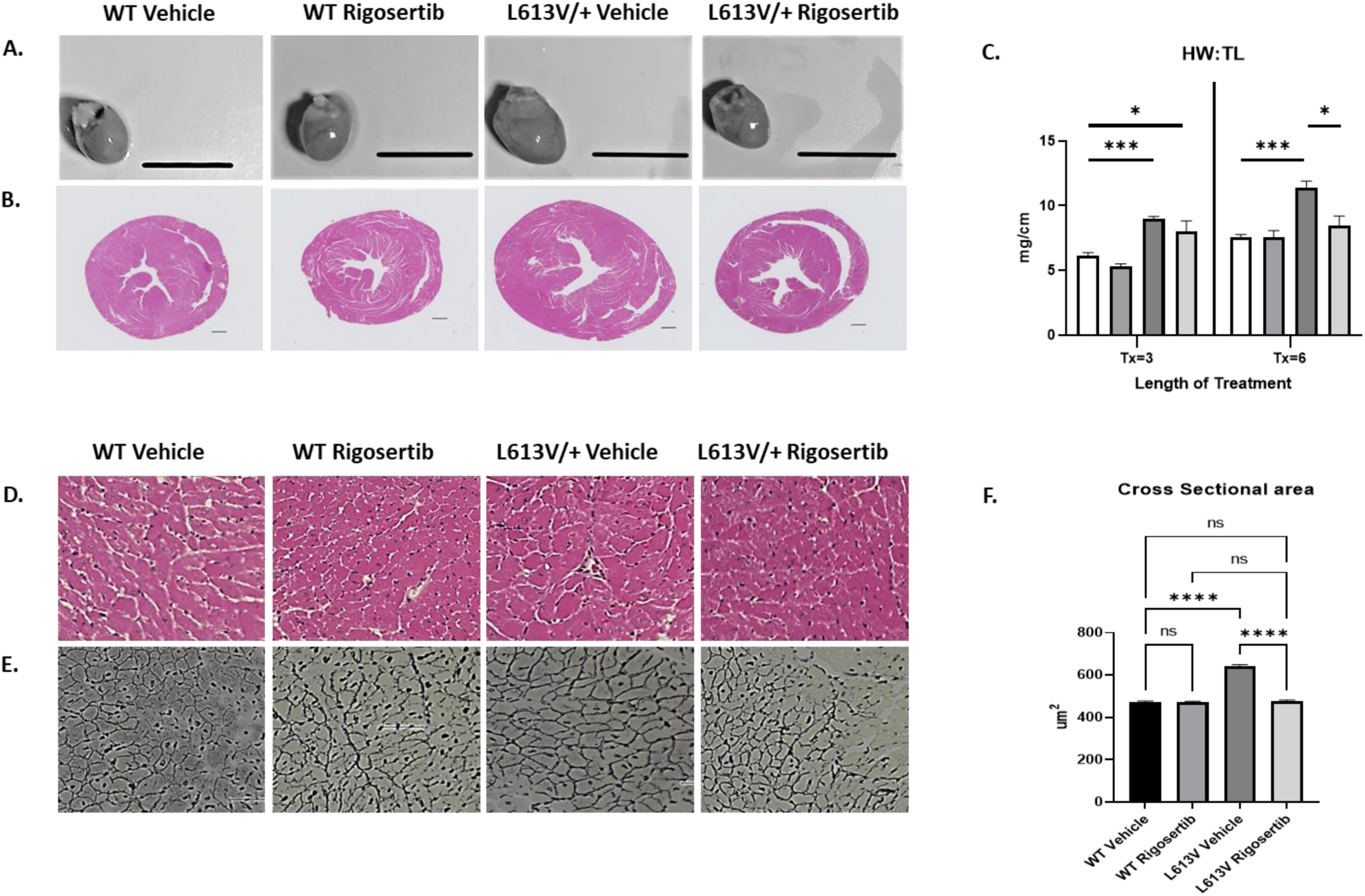
Rigosertib treatment in KI *Raf1^L613V/+^* mice normalizes heart physiology. **A**. Representative photographs of male hearts from 14-week-old WT or RAF1^L613V/+^ mice, either in the presence or absence of intraperitoneally (IP) injected vehicle (water) or rigosertib (100 mg/kg), administered twice a day for a total period of 6 weeks, scale bar=50 mm. **B**. Representative H&E whole heart cross-sections from 14-week-old male WT or RAF1^L613V/+^ mice, either in the presence or absence of intraperitoneally (IP) injected vehicle (water) or rigosertib (100 mg/kg), administered twice a day for a total period of 6 weeks, weeks. Scale bar=1 mm. **C.** Heart weight to tibiae length ratio generated from male 14-week-old WT or RAF1^L613V/+^ mice, either in the presence or absence of intraperitoneally (IP) injected vehicle (water) or rigosertib (100 mg/kg), administered twice a day for a total period of either 3-or 6-weeks. N= 4/group. Data in graphs represent mean ± SEM; statistical significance was determined by 2-way ANOVA with Bonferroni correction, where *p<0.05 and ***p<0.005. **D.** Representative H&E whole heart cross-sections from 14-week-old male WT or RAF1^L613V/+^ mice, either in the presence or absence of intraperitoneally (IP) injected vehicle (water) or rigosertib (100 mg/kg), administered twice a day for a total period of 6 weeks, weeks. Scale bar=50 µm. **E**. Reticulin staining from heart cross-sections of male 14-week-old WT or RAF1^L613V/+^ mice, either in the presence or absence of intraperitoneally (IP) injected vehicle (water) or rigosertib (100 mg/kg), administered twice a day for a total period of 6 weeks, Scale bar= 20 µm. **F**. Frequency distribution of CM area from cross-sections of 14-week-old male WT or RAF1^L613V/+^ mice, either in the presence or absence of intraperitoneally (IP) injected vehicle (water) or rigosertib (100 mg/kg), administered twice a day for a total period of 6 weeks, N=4 mice per group, with at least 1x10^3^ cell counts per heart. Data in graph represents mean ± SEM; statistical significance was determined by 2-way ANOVA with Bonferroni correction, where ****p < 0.0005.

*Raf1^L613V/+^* mice have previously been shown to have increased heart weight (HW) to tibia length (TL) ratios, indicative of their pathological hypertrophy [12]. To determine whether rigosertib normalized heart size, we measured HW:TL ratios in both 3-and 6-week vehicle or rigosertib-treated male and female WT and *Raf1^L613V/+^* mice (**Figure 4C**, **Table 1**, **Table S2**). We observed a progressive improvement in HW:TL ratios in response to rigosertib treatment in both male and female *RAF1^L613V/+^* mice; no changes in HW:TL were observed in WT mice, with or without rigosertib treatment, at either time point (**Figure 4C**, **Table 1**, **Table S2**).

**Table 1.**
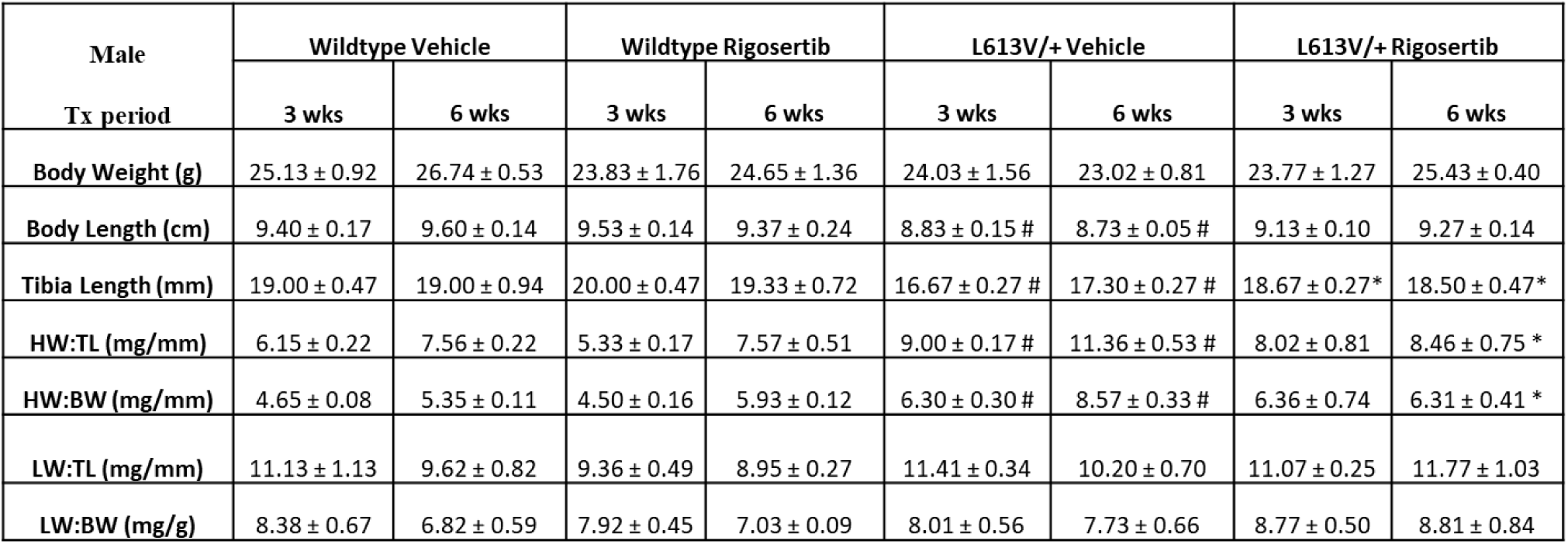
Rigosertib treatment normalized *Raf1^L613V/+^* NS-associated anatomical features. Heart weight (HW), body length, tibia length (TL), HW:TL ratio, HW to body weight (HW:BW) ratio, lung weight (LW) to TL, and LW:BW ratios were all assessed from male WT or *Raf1^L613V/+^* mice that were either intraperitoneally (IP) injected with vehicle (water) or rigosertib (100 mg/kg), administered twice a day, for a total period of 3-weeks or 6-weeks, as indicated, starting at 8 weeks of age. N= 4 mice per group, per time period. Data in tables represent mean ± SEM; statistical significance was determined by 2-way ANOVA with Bonferroni correction, where P<0.05 was deemed significant. Significance is indicated with *, comparison between vehicle-treated *Raf1^L613V/+^* vs. rigosertib-treated *Raf1^L613V/+^* at the same time points; #, comparison between vehicle-treated WT vs. each of the other groups at the same time points, *i.e.* vehicle-treated WT vs. rigosertib-treated WT; vehicle-treated WT vs. vehicle-treated *Raf1^L613V/+^*; vehicle-treated WT vs. rigosertib-treated *Raf1^L613V/+^*.

| Male<br>Tx period | Wildtype Vehicle |  | Wildtype Rigosertib |  | L613V/+ Vehicle |  | L613V/+ Rigosertib |  |
| --- | --- | --- | --- | --- | --- | --- | --- | --- |
|  | 3 wks | 6 wks | 3 wks | 6 wks | 3 wks | 6 wks | 3 wks | 6 wks |
| Body Weight (g) | 25.13 ± 0.92 | 26.74 ± 0.53 | 23.83 ± 1.76 | 24.65 ± 1.36 | 24.03 ± 1.56 | 23.02 ± 0.81 | 23.77 ± 1.27 | 25.43 ± 0.40 |
| Body Length (cm) | 9.40 ± 0.17 | 9.60 ± 0.14 | 9.53 ± 0.14 | 9.37 ± 0.24 | 8.83 ± 0.15 # | 8.73 ± 0.05 # | 9.13 ± 0.10 | 9.27 ± 0.14 |
| Tibia Length (mm) | 19.00 ± 0.47 | 19.00 ± 0.94 | 20.00 ± 0.47 | 19.33 ± 0.72 | 16.67 ± 0.27 # | 17.30 ± 0.27 # | 18.67 ± 0.27* | 18.50 ± 0.47* |
| HW:TL (mg/mm) | 6.15 ± 0.22 | 7.56 ± 0.22 | 5.33 ± 0.17 | 7.57 ± 0.51 | 9.00 ± 0.17 # | 11.36 ± 0.53 # | 8.02 ± 0.81 | 8.46 ± 0.75 * |
| HW:BW (mg/mm) | 4.65 ± 0.08 | 5.35 ± 0.11 | 4.50 ± 0.16 | 5.93 ± 0.12 | 6.30 ± 0.30 # | 8.57 ± 0.33 # | 6.36 ± 0.74 | 6.31 ± 0.41 * |
| LW:TL (mg/mm) | 11.13 ± 1.13 | 9.62 ± 0.82 | 9.36 ± 0.49 | 8.95 ± 0.27 | 11.41 ± 0.34 | 10.20 ± 0.70 | 11.07 ± 0.25 | 11.77 ± 1.03 |
| LW:BW (mg/g) | 8.38 ± 0.67 | 6.82 ± 0.59 | 7.92 ± 0.45 | 7.03 ± 0.09 | 8.01 ± 0.56 | 7.73 ± 0.66 | 8.77 ± 0.50 | 8.81 ± 0.84 |

H&E or reticulin staining of whole heart sections from both male and female *Raf1^L613V/+^* mice showed that the mutant mouse hearts had enlarged individual CMs with increased overall cross-sectional cell surface area. Six-week treatment of *Raf1^L613V/+^* mice with rigosertib normalized these phenotypes to levels similar to WT controls (**Figure 4D-F**; **Figure S1C-E; Table 2A**; **Table S3A**). To confirm these results, we isolated individual CMs from male or female *Raf1^L613V/+^* mice treated with either vehicle or with three or six weeks of rigosertib. We did not observe significant changes in CMs after three weeks of treatment; however, by six weeks, rigosertib normalized length, width, as well as overall cell surface area in both male and female *Raf1^L613V/+^* isolated CMs to levels similar to WT CMs (**Figure 5**; **Figure S2, Table 2B**, **Table S3B**).

**Figure 5.**
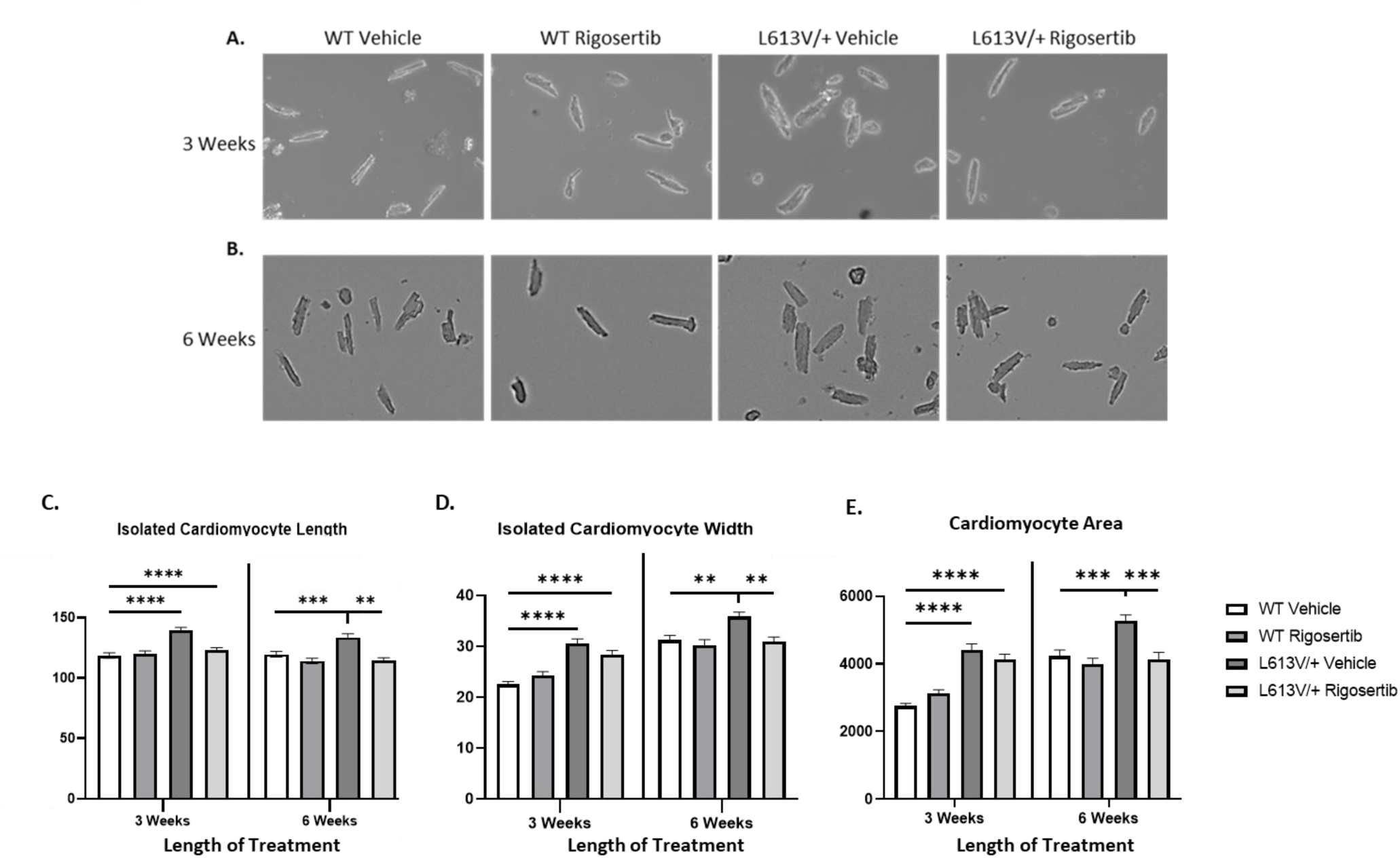
Rigosertib treatment in KI *Raf1^L613V/+^* mice decreases individual CM area, reducing cardiac hypertrophy. Representative photomicrograph of ventricular myocytes isolated from male 14-week-old WT or RAF1^L613V/+^ mice, either in the presence or absence of intraperitoneally (IP) injected vehicle (water) or rigosertib (100 mg/kg), administered twice a day for a total period of either 3-**(A)** or 6-weeks **(B)**. Quantitative assessment of CM length **(C)**, width **(D)**, and total area **(E)**, as assessed from individually isolated CMs isolated from male 14-week-old WT or RAF1^L613V/+^ mice, either in the presence or absence of intraperitoneally (IP) injected vehicle (water) or rigosertib (100 mg/kg), administered twice a day for a total period of either 3-or 6-weeks. N=4 mice per group, with at least 100 cell counts per heart. Data in graphs represent mean ± SEM; statistical significance was determined by 2-way ANOVA with Bonferroni correction, where **p<0.01, ***p<0.005, ****p<0.0005.

**Table 2.** Quantification of CM cross-sectional areas in response to rigosertib treatment in KI *Raf1^L613V/+^* mice. Quantification of cross-sectional CM area (**A**) and measurements of length, width, and total area of individually isolated CMs (**B**) from 14-week-old male WT or RAF1L613V/+ mice, either in the presence or absence of intraperitoneally (IP) injected vehicle (water) or rigosertib (100 mg/kg), administered twice a day for a total period of 3-weeks or 6-weeks, as indicated. N=3 mice per group, with at least 1x10^3^ cell counts per heart. Data in tables represent mean ± SEM; significance is determined by 2-way ANOVA, with Bonferroni correction, where P<0.05 was deemed significant. Significance is indicated with *, comparison between vehicle-treated *Raf1^L613V/+^* vs. rigosertib-treated *Raf1^L613V/+^*at the same time points; #, comparison between vehicle-treated WT vs. each of the other groups at the same time points, *i.e.* vehicle-treated WT vs. rigosertib-treated WT; vehicle-treated WT vs. vehicle-treated *Raf1^L613V/+^*; vehicle-treated WT vs. rigosertib-treated *Raf1^L613V/+^*.

| Male (Tx=6 wks) | Cross-sectional area (um<sup>2</sup>) |
| --- | --- |
| WT Vehicle | 471.36 ± 6.01 |
| WT Treated | 470.13 ± 5.74 |
| L613V Vehicle | 640.35 ± 8.54 # |
| L613V Treated | 476.47 ± 5.19 * |

**Table 2. Quantification of CM cross-sectional areas in response to rigosertib** **treatment in KI *Raf1*<sup>L613V/+</sup> mice.**
| Male | Wildtype Vehicle |  | Wildtype Rigosertib |  | L613V/+ Vehicle |  | L613V/+ Rigosertib |  |
| --- | --- | --- | --- | --- | --- | --- | --- | --- |
| Tx period | 3 wks | 6 wks | 3 wks | 6 wks | 3 wks | 6 wks | 3 wks | 6 wks |
| Length um | 122.3 ± 2.21 | 133.5 ± 3.14 | 129.1 ± 2.58 | 130.9 ± 2.76 | 143.4 ± 3.25 # | 147.7 ± 3.19 # | 144.2 ± 2.96 # | 130.5 ± 3.20 * |
| Width um | 22.53 ± 0.55 | 31.23 ± 0.93 | 24.32 ± 0.68 | 30.26 ± 1.06 | 30.52 ± 0.96 # | 35.86 ± 0.89 # | 28.4 ± 0.83 # | 30.9 ± 0.91 * |
| Area um<sup>2</sup> | 2753 ± 78.65 | 4226 ± 182.00 | 3126 ± 105.80 | 3986 ± 182.80 | 4409 ± 183.20 # | 5277 ± 168.20 # | 4133 ± 153.30 # | 4138 ± 197.70 * |

### Rigosertib treatment of KI Raf1^L613V/+^ mice normalized cardiac function

To determine the effects of rigosertib on cardiac function, we conducted echocardiographic analysis in these mice and found that the vehicle-treated *Raf1^L613V/+^* mice showed significant progressive left ventricular hypertrophy at both 11-and 14-weeks of age, as indicated by increased LV mass as well as thickened IVS and LVPW as compared with WT controls (**Figure 6**, **Table 3**, **Table S4**). Treatment of *Raf1^L613V/+^* mice with rigosertib, however, normalized the hypertrophy phenotype by the end of the 6-week treatment period (**Figure 6**, **Table 3**, **Table S4**). Interestingly, we also observed increased fractional shortening (FS) and ejection fraction (EF) in the *Raf1^L613V/+^* mutant mice, consistent with HCM, which was reduced in the rigosertib-treated mice (**Table 3**, **Table S4**), suggesting overall cardiac functional normalization. No functional changes or effects were observed in WT hearts over the 3-or 6-week treatment period, either with or without rigosertib (**Table 3**, **Table S4**).

**Figure 6.**
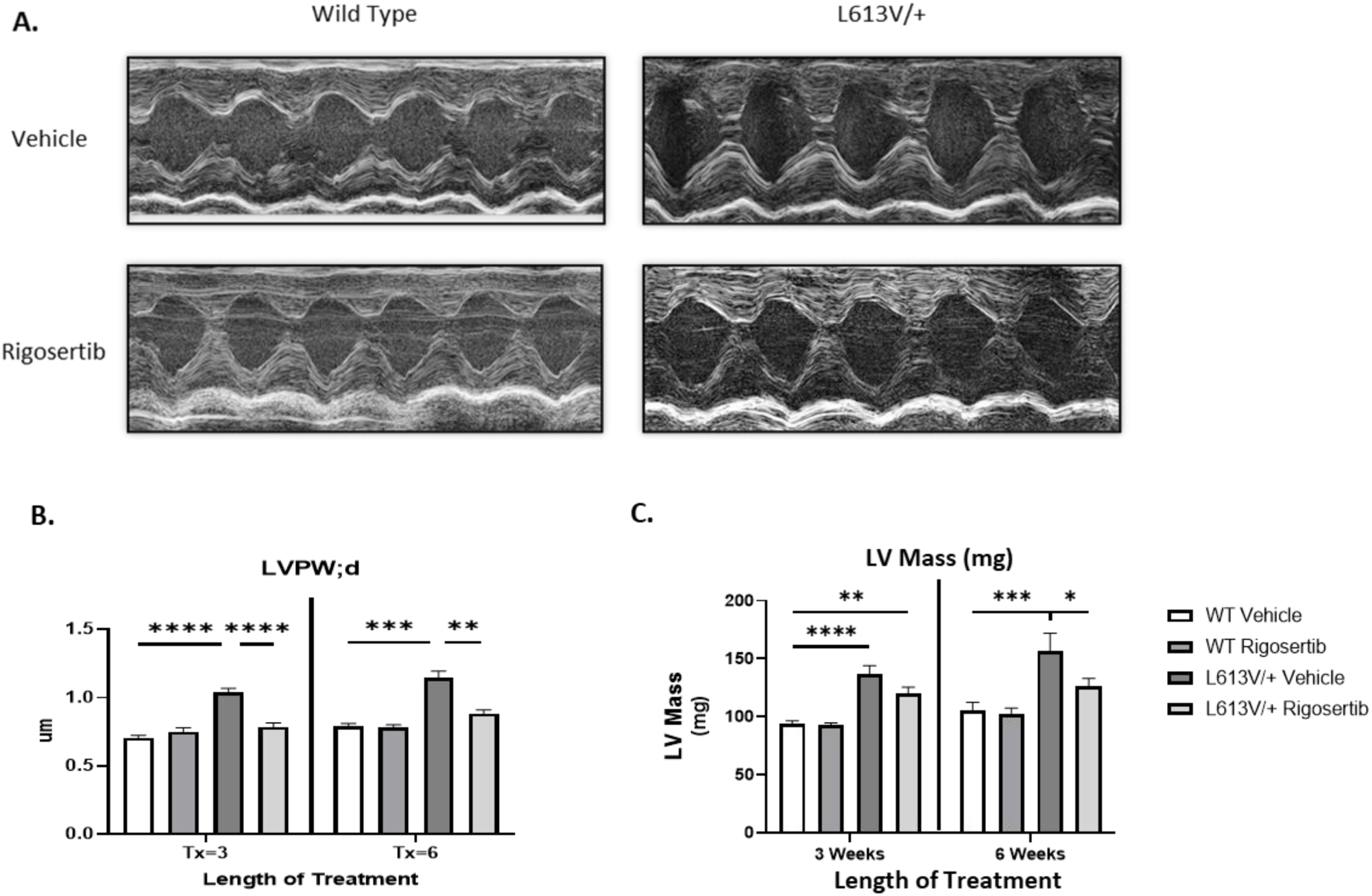
Rigosertib treatment in KI *Raf1^L613V/+^* mice normalizes cardiac function. **A**. Representative echocardiography of male 14-week-old WT or RAF1^L613V/+^ mice, either in the presence or absence of intraperitoneally (IP) injected vehicle (water) or rigosertib (100 mg/kg), administered twice a day for a total period of 6-weeks. Quantification of **B.** LVPWd and **C.** LV mass (n=4/group). LVPW;d=left ventricular posterior wall thickness in diastole. Data in graphs represent mean ± SEM; statistical significance was determined by 2-way ANOVA with Bonferroni correction, where *p<0.05, **p<0.01, ***p<0.001, ****p<0.0001.

**Table 3.** Echocardiographic analysis of WT vs *Raf1^L613V/+^* mice. WT or *Raf1^L613V/+^*male mice were either treated with intraperitoneally (IP) injected vehicle (water) or rigosertib (100 mg/kg), administered twice a day, for a total period of 3-weeks or 6-weeks, as indicated, starting at 8 weeks of age. Echocardiographic measurements in M-mode to measure cardiac function were taken at both 3-and 6-week time points. Tx=treatment; BPM=beats per minute; diameter;d= diameter in diastole; SV= stroke volume, EF= ejection fraction, FS= fractional shortening, LV= Left Ventricular; LVAW;d= left ventricular anterior wall thickness in diastole; LVPW;d= left ventricular posterior wall thickness in diastole. N=4 mice per group, per time point. Data in tables represent mean ± SEM; statistical significance was determined by 2-way ANOVA with Bonferroni correction, where P<0.05 was deemed significant. Significance is indicated with *, comparison between vehicle-treated *Raf1^L613V/+^* vs. rigosertib-treated *Raf1^L613V/+^* at the same time points; #, comparison between vehicle-treated WT vs. each of the other groups at the same time points, *i.e.* vehicle-treated WT vs. rigosertib-treated WT; vehicle-treated WT vs. vehicle-treated *Raf1^L613V/+^*; vehicle-treated WT vs. rigosertib-treated *Raf1^L613V/+^*. Note: no differences in WT groups were noted at either time point, either in the presence or absence of rigosertib, suggesting no adverse effects of the inhibitor in normal hearts.

| Echo Data (Male) | Wildtype Vehicle |  | Wildtype Rigosertib |  | L613V/+ Vehicle |  | L613V/+ Rigosertib |  |
| --- | --- | --- | --- | --- | --- | --- | --- | --- |
| Tx period | 3 wks | 6 wks | 3 wks | 6 wks | 3 wks | 6 wks | 3 wks | 6 wks |
| Heart Rate (BPM) | 665.4 ± 26.45 | 680.59 ± 16.26 | 629.2 ± 22.46 | 641.67 ± 32.56 | 624.2 ± 15.96 | 691.88 ± 18.69 | 613.7 ± 18.4 | 661.19 ± 22.30 |
| Diameter;d (mm) | 3.828 ± 0.11 | 3.76 ± 0.11 | 3.688 ± 0.08 | 3.77 ± 0.09 | 3.602 ± 0.09 | 3.54 ± 0.17 | 4.16 ± 0.12 * | 3.86 ± 0.11 |
| SV (uL) | 43.86 ± 2.46 | 44.38 ± 2.57 | 43.54 ± 1.62 | 45.24 ± 1.77 | 46.92 ± 3.16 | 42.74 ± 3.83 | 56.28 ± 2.78 * # | 49.44 ± 2.96 |
| EF (%) | 72.76 ± 2.13 | 73.35 ± 1.11 | 76.76 ± 1.23 | 74.54 ± 1.54 | 82.19 ± 1.46 # | 81.6 ± 2.72 # | 74.04 ± 1.54 * | 76.35 ± 0.89 |
| FS (%) | 38.88 ± 1.91 | 41.82 ± 0.91 | 43.91 ± 1.20 | 42.95 ± 1.39 | 53.7 ± 1.43 # | 50.03 ± 2.84 # | 42.46 ± 1.82 * | 44.66 ± 0.71 |
| LV Mass (mg) | 93.57 ± 2.89 | 104.87 ± 7.42 | 92.96 ± 1.70 | 102.04 ± 5.39 | 136.5 ± 7.60 # | 156.46 ± 15.51 # | 120.1 ± 5.26 # | 126.28 ± 6.56 * |
| IVS;d (mg) | 0.7221 ± 0.02 | 0.78 ± 0.02 | 0.7352 ± 0.01 | 0.76 ± 0.01 | 0.9637 ± 0.06 # | 1.08 ± 0.05 # | 0.767 ± 0.03 * | 0.88 ± 0.03 * |
| LVPW;d (mm) | 0.7048 ± 0.02 | 0.79 ± 0.02 | 0.7489 ± 0.03 | 0.78 ± 0.02 | 1.038 ± 0.03 # | 1.144 ± 0.05 # | 0.784 ± 0.03 * | 0.88 ± 0.03 * |

Of note, rigosertib significantly increased the LVIDd of the male *Raf1^L613V/+^* mice at three weeks of treatment; however, this effect appeared transient, as the diameter dimension normalized to that of WT controls by six weeks of rigosertib treatment (**Table 3**). In contrast, three weeks of rigosertib treatment was sufficient to normalize LV mass in the female *Raf1^L613V/+^* mice, (**Table S4 vs. Table 2**), a sexual dimorphism consistent with the fact that HCM is generally milder in females than in males with the *Raf1^L613V/+^* genotype.

### Rigosertib treatment of KI Raf1^L613V/+^ mice reduced fetal gene expression, suggesting reversal of pathological hypertrophy

Given the strong phenotypic and functional improvements following rigosertib treatment of *Raf1^L613V/+^*mutant mice, we next examined its effects on the molecular signaling of treated hearts. To assess this, we first conducted quantitative PCR (qPCR) analysis to measure changes in fetal genes, including atrial natriuretic factor (*Anf*), brain natriuretic peptide (*Bnp*), alpha myosin heavy chain (*Myh6*), and beta myosin heavy chain (*Myh7*) (**Figure S3A**). While the mRNA levels for *Myh6* were decreased in *Raf1^L613V/+^* mouse hearts, they were normalized to levels similar to WT in *Raf1^L613V/+^* rigosertib-treated hearts. Consequently, the ratio of *Myh7* to *Myh6* was also increased, indicating improvement in overall cardiac function in response to rigosertib. In addition, we observed a trend towards normalized fetal gene expression of both *Anp* and *Bnp* in response to rigosertib treatment of *Raf1^L613V/+^* mice, though significance was not reached (**Figure S3B-C**).

### Rigosertib treatment of KI Raf1^L613V/+^ mice reduced hypertrophy-induced activation of Erk and Akt

Rigosertib has previously been shown to be a dual PI3K/AKT and ERK/MAPK pathway inhibitor, both of which can be altered in RASopathy patients. To assess the effects of these pathways in rigosertib-treated hearts, we isolated whole hearts from either WT or *RAF1^L613V/+^* mice treated with vehicle (water) or rigosertib (100 mg/kg). Drug was administered twice a day, beginning at 8 weeks of age, for a period of six weeks. While treatment of rigosertib trended towards reducing Akt activation in lysates from *Raf1^L613V/+^*hearts vs. vehicle treatment, the results did not reach significance (**Figure S3D-E**); rigosertib also did not have any effects on Akt activation in WT hearts (**Figure S3D-E**). Similarly, no changes in Erk activation were detected in whole heart lysates isolated from either WT or *Raf1^L613V/+^* mice, with or without rigosertib treatment (**Figure S3D, F**).

One possible explanation for these results could be that treatment with rigosertib over 6 weeks could normalize any hyperactivation of Erk and Akt, making acute changes in signaling difficult to observe. Alternatively, other cell types in whole heart could also perhaps mask effects of rigosertib-induced inhibition of these pathways in CMs. Therefore, to more fully elucidate the signaling effects of rigosertib directly in CMs, we isolated primary CMs from WT and *Raf1^L613V/+^* mutant mice, treated them with rigosertib for 12 hours *in vitro*, and then subsequently stimulated them with either vehicle, AngII (20 ng/ml) or IGF (50 ng/ml) for 5 minutes, to determine acute effects of drug treatment on Erk and Akt signaling, respectively. At baseline, we did not observe any significant changes in either Erk or Akt activity, consistent with previous findings [12]. In response to treatment with AngII, an upstream activator of the ERK/MAPK signaling cascade, Erk1/2 phosphorylation was significantly increased in *Raf1^L613V/+^* CMs as compared to WT CMs (**Figure 7A-B**). Similarly, in response to IGF stimulation, a mediator of PI3K/AKT signaling, *Raf1^L613V/+^*CMs exhibited significantly increased Akt phosphorylation as compared to WT controls (**Figure 7A, 7C**). Rigosertib treatment of mutant cardiomyocytes, however, was able to normalize the Erk and Akt activities of *Raf1*-mutant CMs to levels similar to those of WT (**Figure 7A-C**).

**Figure 7.**
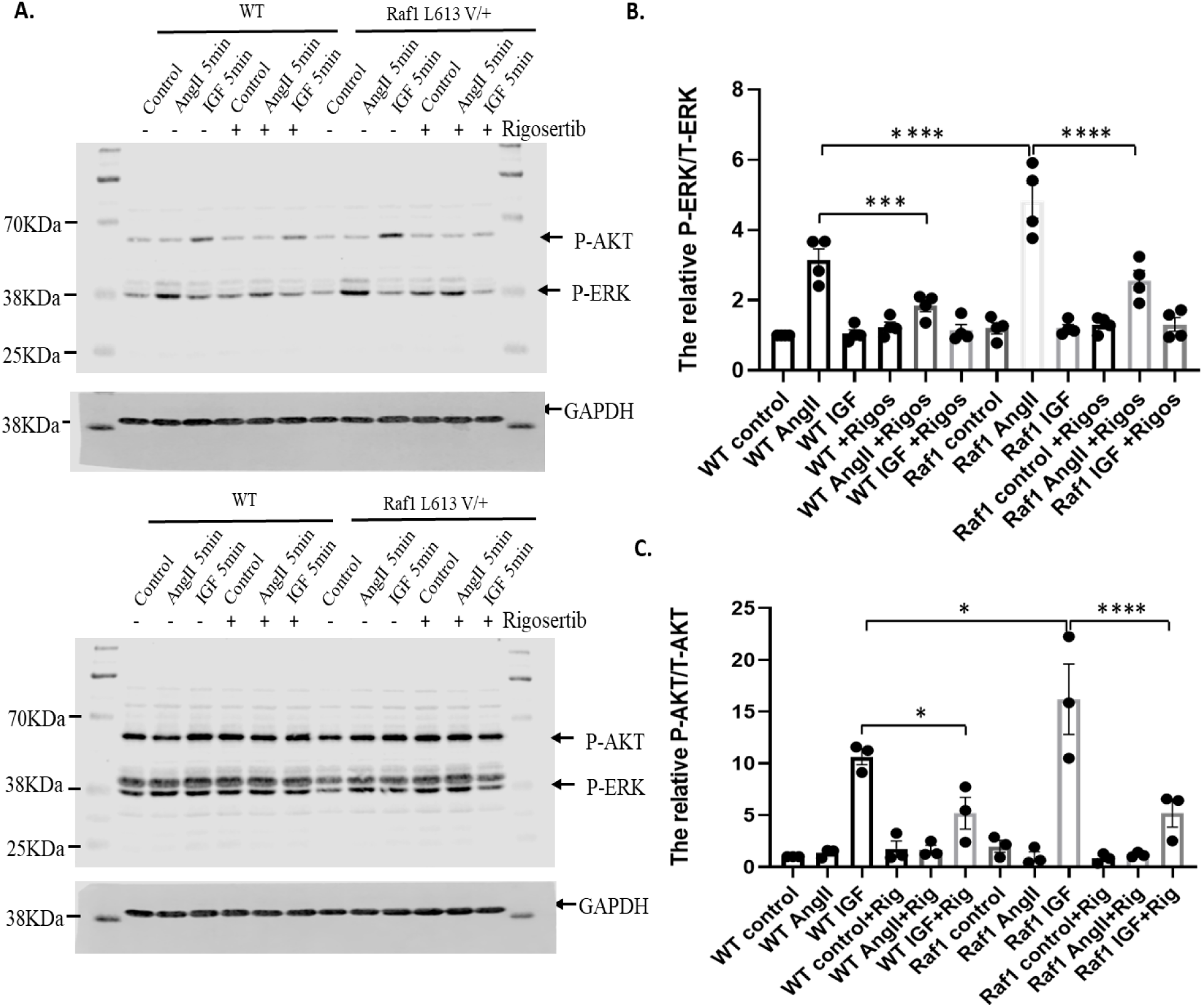
Rigosertib treatment in KI *Raf1^L613V/+^* mice inhibits hyperactivation of ERK and AKT activities in response to AngII and IGF stimulation, respectively. Representative immunoblots of mouse heart lysates isolated from WT or RAF1^L613V/+^ mice littermates, cultured for 12 hours in the absence or presence of rigosertib, and then subjected to 5 minutes of stimulation with either vehicle, AngII (20 ng/ml) or IGF (50 ng/ml). The heart lysates were immunoblotted with phospho-ERK1/2 or phospho-AKT, as indicated, followed by total ERK1/2 and total AKT to control for total protein expression. Experimental samples were also immunoblotted with anti-GAPDH to normalize loading within each gel. Quantification of **B.** p-ERK/total ERK and **C.** p-AKT/total AKT immunoblotting data from n= 3 experimental replicates are indicated. Data represent mean ± SEM; statistical significance was determined by 2-way ANOVA with Bonferroni correction, where *p<0.05; ***p<0.001; ****p<0.0001.

### Rigosertib reduces craniofacial abnormalities and induces linear growth in Raf1^L613V/+^ mice

In addition to HCM, previous work has shown that *Raf1^L613V/+^* mice display several of the other characteristic NS phenotypes, including craniofacial abnormalities, reduced stature, and shortened tibias [12]. To determine whether rigosertib affects non-cardiac NS-associated phenotypes, we conducted μCT scans of the craniofacial areas and showed that treatment with rigosertib normalized craniofacial inner canthal width in the *Raf1^L613V/+^* mice to measures similar to those in WT (**Figure 8A-B**, **Table S5**). Skull lengths, widths, and nose lengths from treated *Raf1^L613V/+^* mice also trended towards normalized WT measures but did not reach significance in our study (**Table S5**).

**Figure 8.**
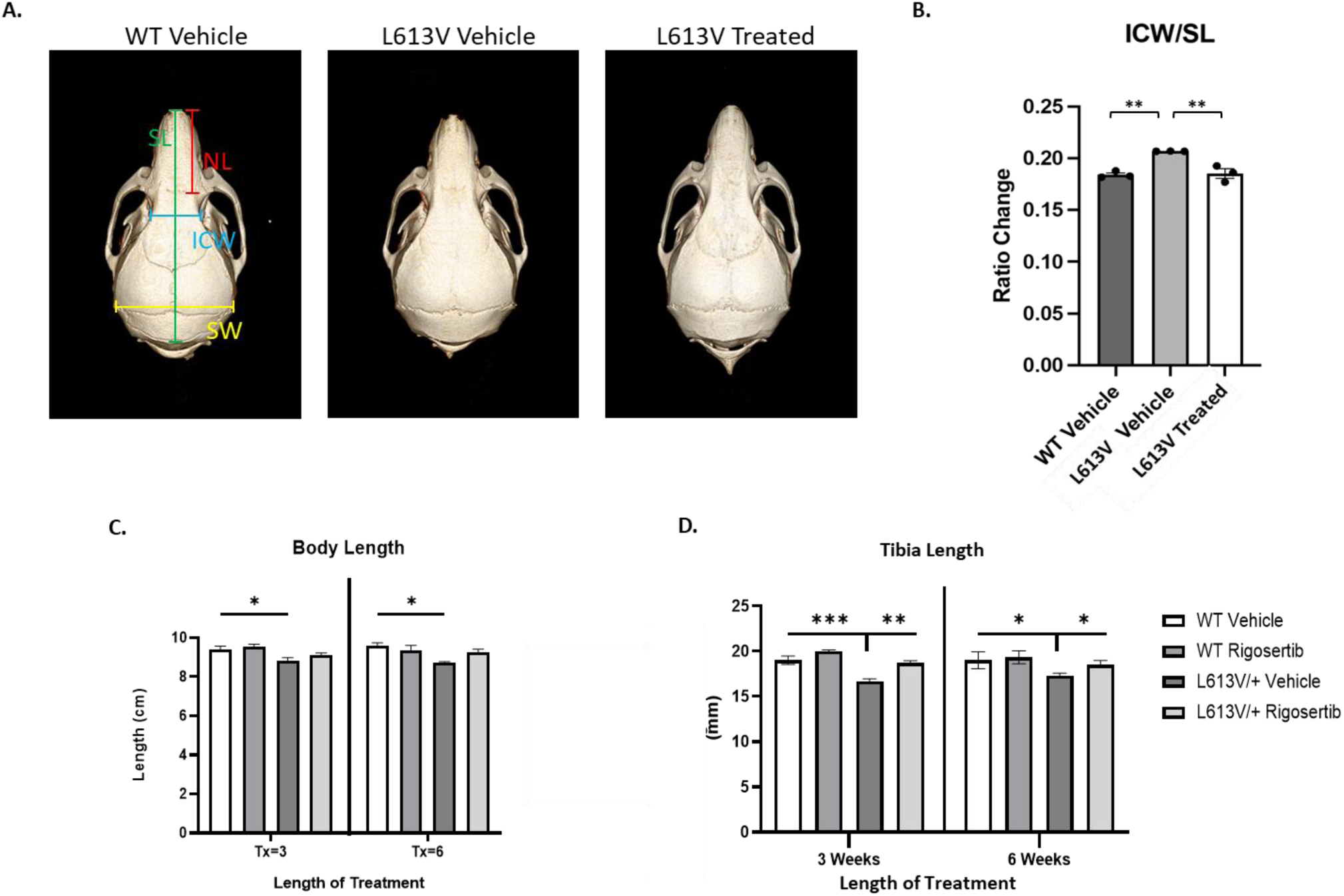
Rigosertib functions to ameliorate multiple NS syndromic features in RAF1^L613V/+^ mice. **A.** Representative μCT scans showing measures of skull lengths, widths, inner canthal width, and nose lengths; Quantification of the inner canthal width (**B**), body length (**C**), and tibia length (**D**) from male 14-week-old WT or RAF1^L613V/+^ mice, either in the presence or absence of intraperitoneally (IP) injected vehicle (water) or rigosertib (100 mg/kg), administered twice a day for a total period of either 3-or 6-weeks. N= 4/group. Data in graphs represent mean ± SEM; statistical significance was determined by 2-way ANOVA with Bonferroni correction, where *p<0.05, **p<0.01, and ***p<0.005.

In addition to assessing changes in facial dysmorphia, we also measured total body length and tibia length in both 3-and 6-week treated male and female *Raf1^L613V/+^* mice (**Figure 8C-D**, **Table 1**, **Table S2**). Rigosertib treatment led to a striking reversal of the short stature phenotype and normalized tibia lengths in both male and female rigosertib-treated *Raf1^L613V/+^* mice (**Figure 8C-D**, **Table 1**, **Table S2**). This indicates that rigosertib treatment not only ameliorates the HCM phenotype in *RAF1*-assoicated NS but can also lead to positive benefits on non-cardiac aspects of this disorder as well.

## Discussion

Treatments for RASopathies remain limited. Our data indicate that rigosertib, a compound currently in multiple clinical trials and with demonstrated low toxicity [25], is a strong candidate for the treatment of RASopathy-associated HCM. Treatment with rigosertib led to an increase in survival of multiple transgenic RASopathy Drosophila models. We also observed an improvement in adult heart function and a rescue of the hypertrophic phenotype in cardiac-specific *RAF1^L613V^ Drosophila* that were fed rigosertib. Further, rigosertib treatment at concentrations as low as 100 nM both prevented and reversed the hypertrophic phenotype in human iPSC-derived *RAF1^S257L^* CMs. These findings were validated in *Raf1^L613V/+^* mice, where we showed that rigosertib treatment reversed HCM, reduced CM size, improved cardiac function, and normalized cardiac pathology. Further and important for clinical considerations, rigosertib had significant positive effects on other NS-associated pathologies, including craniofacial anomalies, short stature and tibia length. Overall, our data suggest rigosertib is a promising treatment for *RAF1*-associated HCM and, perhaps, other RASopathies and/or RASopathies-associated pathologies as well.

Rigosertib was initially developed, alongside several other styryl benzyl sulfones, to address elevated cyclin activity that leads to tumorigenesis as a consequence of elevated Polo-like kinase 1 (PLK1) [20]. The treatment showed promise early on, as the drug-induced cytotoxicity in tumor cells while leaving healthy cells unharmed [25]. Having both a high level of efficacy against a multitude of tumor types and a favorable toxicity profile [38], rigosertib has been a compound of great interest for treatment of disease. Interestingly, the mechanism for its regulation was later found to not involve binding to PLK1 directly; instead, rigosertib functioned as a non-ATP competitive inhibitor, essentially acting as a RAS binding domain mimetic [39], inhibiting multiple kinase pathways downstream of RAS, including PI3K/AKT and RAS/MAPK signaling [20]. Recently, rigosertib has been shown to modulate the PLK-1 pathway in squamous cell carcinoma in the setting of recessive dystrophic epidermolysis bullosa (RDEB-SCC) [27], triple negative breast cancer [38, 40] and hepatitis C virus proliferation [41].

Increased ERK and MEK activities have been previously demonstrated in *RAF1^L613V/+^* mouse hearts in response to transverse aortic constriction (TAC), a chronic pathological stressor that leads to cardiac hypertrophy and heart failure [12]. Interestingly, treatment with the MEK inhibitor mirdametinib normalized these cardiac defects, suggesting a role for aberrant ERK signaling in *RAF1*-associated HCM development [12]. In addition, we previously showed that *RAF1*-mutant iPSC-derived CMs mediate HCM through hyperactivation of MEK1/2, but not ERK1/2, mediating myofibrillar disarray [11]. Surprisingly, in this case, we found that the enlarged CM phenotype occurred as a consequence of increased extracellular regulated kinase 5 (ERK5) signaling, a pathway not previously known to be involved in NS [11]. Additional studies are required to assess whether rigosertib may affect ERK5 and/or other parallel signaling pathways in RASopathies associated with HCM.

Rigosertib is a dual RAS/MAPK and PI3K/AKT inhibitor and herein displays a positive effect on reversing HCM in *RAF1*-associated RASopathy. Indeed, several studies similarly suggest positive effects of RAS/MAPK and PI3K/AKT inhibition in the heart. For example, prior pharmacological studies have shown that treatment of CMs with an AKT inhibitor prevents hypertrophy evoked by stimulation by various agonists [42–45]. Inhibition of AKT/mTOR also attenuates or reverses pressure overload-associated cardiac hypertrophy [43, 46, 47]. In RASopathies, we generated an NSML KI mouse model of the *PTPN11* mutation Y279C (*Ptpn11^Y279C/+^)* that recapitulated the human disorder, with short stature, craniofacial dysmorphia, and HCM; interestingly, these mice showed aberrant agonist-evoked ERK/MAPK signaling, but increased basal and agonist-induced AKT/mTOR activity [8]. Moreover, in these mice, the HCM-associated cardiac defects were completely reversed with treatment by rapamycin, an inhibitor of mTOR [8]. Similarly, NSML iPSC-derived CMs demonstrated higher sarcomeric disorganization and increased cell size, indicative of cardiac hypertrophy [48]. Use of a specific AKT inhibitor, ARQ 092, in another NSML iPSC-derived CM line blocked pathological HCM [48]. Together, these data support the notion that PI3K/AKT signaling plays a regulatory role in the development of pathological HCM, and that inhibition of this pathway may positively affect and reverse pathological cardiac outcomes.

As with PI3K/AKT signaling, preclinical studies have also suggested that chronic inhibition of RAS/MAPK signaling may have beneficial and cardioprotective effects against development of cardiac hypertrophy and heart failure [49, 50]. Blocking RAS/MAPK signaling also leads to decreased cardiac fibrosis in response to pathological stress, including ischemia/reperfusion (I/R) injury [51, 52]. Indeed, MEK inhibitors have recently gained traction in RASopathies. Specifically, in 2020, the FDA approved selumetinib to treat children with NF1 with symptomatic, inoperable plexiform neurofibromas and low-grade gliomas (LGGs); the phase 2 trial of selumetinib for treatment of LGGs resulted in 40% partial response and 96% of patients with 2 years of progression-free survival [53]. More recently, mirdametinib, an oral allosteric MEK inhibitor, is being evaluated for treatment of NF1 patients [54]. While these studies appear promising, at least one study suggests that long-term inhibition of RAS/MAPK signaling may elicit adverse cardiovascular outcomes in time, through increased activation of cAMP and PKA [55].

In summary, our data identify rigosertib as a potent therapeutic candidate that can reverse *RAF1*-associated HCM. Although these findings hold promise for the treatment of hypertrophy in this and other RASopathies, additional experiments, including optimization of dose and treatment duration as well as long-term efficacy and toxicity will be needed to fully assess the clinical implications for its use in patients with these disorders. This is especially true given RAS/MAPK and PI3K/AKT pathways are critically involved in important biological processes that regulate growth and differentiation. In all, we find rigosertib to be a highly promising, well-tolerated, and highly effective therapeutic for treatment of RAF1-associated HCM and, potentially, other RASopathies-dependent pathologies as well.

## Non-standard Abbreviations and Acronyms

ANF: Atrial natriuretic factor
BNP: Brain natriuretic peptide
bp: Base pair
CHD: Congenital heart disease
CFC: Cardiofaciocutaneous syndrome
CM: Cardiomyocyte
CRISPR: Clustered regularly interspaced short palindromic repeats
CS: Costello syndrome
cTNT: Cardiac troponin T
DMSO: Dimethyl sulfoxide
EBs: Embryoid bodies
EF: Ejection fraction
ERK: Extracellular signal regulated kinase
FACS: Fluorescence-activated cell sorting
FLCs: Fibroblast-like cells
FS: Fractional shortening
HCM: Hypertrophic cardiomyopathy
H&E: Hematoxylin/eosin
HW: Heart weight
IP: Intraperitoneally
iPSCs: Inducible pluripotent stem cells
I/R: ischemia/reperfusion
KI: Knock-in
LVIDd: Left ventricular internal diameter end-diastole
LVIDs: Left ventricular internal diameter end-systole
LVPW: Left ventricular posterior wall thickness
MAPK: Mitogen-activated protein kinase
MEF: Mouse embryonic fibroblast
MEK: Mitogen-activated protein kinase kinase
MTG: Monothioglycerol
MMRRC: Mutant Mouse Resource & Research Centers
MYH6: α-myosin heavy chain
MYH7: β-myosin heavy chain
NF1: Neurofibromatosis 1
NS: Noonan syndrome
NSML: Noonan syndrome with multiple lentigines
PI3K: Phosphatidylinositol 3-kinase
RIPA: Radioimmunoprecipitation
*Raf1^L613V/+^*: RASopathy mice heterozygous for the *Raf1* L613V allele
Rigosertib: 2-[2-methoxy-5-[[(E)-2-(2,4,6-trimethoxyphenyl) ethenyl] sulfonylmethyl] anilino] acetic acid (ON01910)
RFLP: Restriction-fragment length polymorphism
SEM: Standard error of the mean
SLN: Sarcolipin
SDS-PAGE: Sodium dodecyl-sulfate polyacrylamide gel electrophoresis
ssODN: Single-stranded oligonucleotide donor
SV: Stroke volume
*tin>hRaf1L613V*: Cardiac-specific tin-GAL4 driving UAS-*hRaf1* L613V transgene
TL: Tibia length
WT: Wildtype

## Acknowledgments

We would like to thank Onconova Therapeutics (now Traws Pharma) for providing us with the rigosertib compound and for their consultation and discussion of these data.

